# Structural insight into the nuclear transportation mechanism of PPARγ by Transportin-1

**DOI:** 10.1101/2024.09.17.612794

**Authors:** Sachiko Toma-Fukai, Yutaro Nakamura, Akihiro Kawamoto, Hikaru Shimizu, Koki Hayama, Ruri Kojima, Kanami Yoshimura, Masaki Ishii, Mika Hirose, Toshiaki Teratani, Shinya Ohata, Takayuki Kato, Hironari Kamikubo, Toshimasa Itoh, Kengo Tomita, Toshiyuki Shimizu

**Affiliations:** Graduate School of Pharmaceutical Sciences, The University of Tokyo, 7-3-1 Hongo, Bunkyo-ku, Tokyo 113-0033, Japan; Graduate School of Science and Technology, Nara Institute of Science and Technology, 8916-5 Takayama-cho, Ikoma, Nara 630-0192, Japan; Institute for Protein Research, Osaka University, Suita, Osaka, 565-0871, Japan; Japan Science and Technology Agency, PRESTO, Kawaguchi, Japan; Research Institute of Pharmaceutical Sciences, Faculty of Pharmacy, Musashino University,Tokyo 202-8585, Japan; Division of Gastroenterology and Hepatology, Department of Internal Medicine, Keio University School of Medicine, 35 Shinanomachi, Shinjuku-ku, Tokyo 160-8582, Japan; Laboratory of Drug Design and Medicinal Chemistry, Showa Pharmaceutical University, 3-3165 Higashi-Tamagawagakuen, Machida, Tokyo 194-8543, Japan

**Keywords:** PPARγ, Transporin-1, nuclear transportation, single particle Cryo-EM

## Abstract

The spatial and temporal control of protein is essential for normal cellular function. Proteins working in the nucleus have nuclear localization signal (NLS) sequences and are escorted into the nucleus by cognate nuclear transport receptors. A wealth of experimental data about NLS has been accumulated, and nuclear transportation mechanisms are established at the biochemical and structural levels.

The peroxisome proliferator-activated receptors (PPARs) are ligand-dependent transcription factors that control various biological responses. We recently reported that the transportation of PPARγ is mediated by Transportin-1, but PPARγ lacks a typical NLS sequence recognized by Transportin-1. Moreover, the recognition mechanism remains largely unknown.

In this study, we determined the Cryo-EM structure of PPARγ in complex with Transportin-1 and revealed that Transportin-1 gripped the folded DNA binding domain and the Hinge region of PPARγ, indicating that PPARγ recognizes a folded domain with an extended region as a nuclear localization signal, not a canonical unstructured signal sequence, confirmed by the mutation analyses in vitro and in cultured cells. Our study is the first snapshot structure working in nuclear transportation, not in transcription, of PPARγ.

## Introduction

The spatial and temporal control of protein is essential for normal cellular function. The dysfunction of the regulation causes disease(*1*). One of the most famous spatial regulation processes is the nuclear transportation of proteins. Nuclear proteins should enter the nucleus after translation in the cytosol.

The peroxisome proliferator-activated receptors (PPARs) are a member of the nuclear receptor family proteins considered an integral part of essential human drug target families (GPCR, ion channels, protein kinase, and nuclear receptor)(*2*). PPARs control cellular processes, including lipid regulation and carbohydrate metabolism. The proteins sense lipid ligands (steroid and thyroid hormones, vitamins, lipid metabolites, and xenobiotics) and activate target genes(*3, 4*). PPARs consist of three subtypes: PPARα, PPARβ (also known as PPARδ), and PPARγ. The family proteins commonly harbor the N-terminal domain, DNA binding domain (DBD), the hinge region, and the ligand binding domain (LBD)(*3*) (*SI Appendix*, Fig. S1). The binding of PPAR to specific DNA sequences requires heterodimerization with a second member of the nuclear receptor family, the retinoic X receptor (RXR)(*3, 5*). The binding of agonist ligands to PPAR triggers a conformation change that recruits transcriptional coactivators, including members of the steroid receptor coactivator (SRC) family(*6*).

PPARγ, the best-studied member of the PPAR family, is expressed in white and brown adipocytes and regulates adipocyte differentiation, lipid storage, and release. PPARγ is the master regulator of fat-cell formation(*5*). One class of PPARγ ligands, the thiazolidinediones, which includes the drug rosiglitazone, are effective insulin sensitizers and have been shown to improve glucose uptake and lower hyperglycemia and hyperinsulinaemia(*7-10*). The PPARs are potential therapeutic targets for atherosclerosis, inflammation, and hypertension(*11*).

In contrast to the enormous studies about gene activation mechanisms, the molecular mechanism of the nuclear localization of the PPAR family remains elusive. Nuclear transportation receptors, known as Karyopherin-β2 (Kap) family proteins, transport cargo proteins into/from nuclear by shuttling between nuclear and cytosol. Kap proteins recognize intrinsically disordered linear elements known as nuclear localization or export signals (NLSs or NESs) and/or folded domains of cargos, while the recognition mechanisms of the cargo proteins without canonical NLS/NES by Kaps are not clearly understood(*1*).

Recently, our group reported that Transportin-1 (Trn1), also known as karyopherin-β2 (Kapβ2), works as the nuclear transportation receptor of PPARγ in a redox-dependent manner(*12*). Trn1 is a member of the Kapβ family proteins and directly recognizes the PY-NLS sequence and the RGG regions of the cargo proteins(*13-18*). However, PPARγ lacks the above the NLS sequence and its recognition mechanism remains largely unknown.

In this study, we determined the Cryo-EM structure of the Trn1 in complex with the almost full-length PPARγ, including three functional regions (DBD, hinge region, and LBD), and revealed that Trn1 enfolds the DBD and the hinge region of PPARγ on the concave surface. Biochemical experiments in vitro and nuclear transportation assay in cultured cells confirmed that the DBD and N-terminal domain of the hinge region work as a nuclear localization signal structure. Our study is the first snapshot structure working in nuclear transportation, not in transcription, of PPARγ.

## Results

### Trn1 exhibits binding affinity for the DNA binding domain and its flanking hinge region of PPARγ

We prepared GST-PPARγ (DBD) and GST-PPARγ (102-505) containing DBD, hinge region, and LBD, hereafter referred to as PPARγ (DHL), to identify the Trn1 binding region of PPARγ. The GST-pulldown assays showed that DBD can bind to full-length Trn1 but the binding affinity of GST-PPARγ (DBD) was weaker than that of GST-PPARg (DHL) (Fig. 1*A*, *SI Appendix*, Fig. S2). ITC analysis showed that the dissociation constant (*K*_D_) of PPARg (DHL) to Trn1 was 200 nM (*SI Appendix*, Fig. S3). Next, to evaluate the solution property of the complex, we performed the SEC-MALS and SEC-SAXS analyses using full length Trn1 and Trn1δloop that is a mutant in which the HEAT repeat 8 loop is replaced with the GGSGGSG linker (*SI Appendix*, Fig. S2)(*15*). The SEC-MALS analysis showed that both full-length Trn1 and Trn1Δloop binds to PPARγ (DBD) or PPARγ (DHL) with the 1:1 molar ratio (*SI Appendix*, Fig. S4). SEC-SAXS solution structures of the complex showed that the concave face of Trn1 molecule was filled with the dummy model, strongly suggesting that Trn1 grips PPARγ using the concave face of Trn1 in the solution (*SI Appendix*, Fig. S5). No apparent structural difference in a binding manner between Trn1 and Trn1Δloop suggests that the HEAT repeat 8 loop does not affect the binding mode of PPARγ.

**Figure 1.**
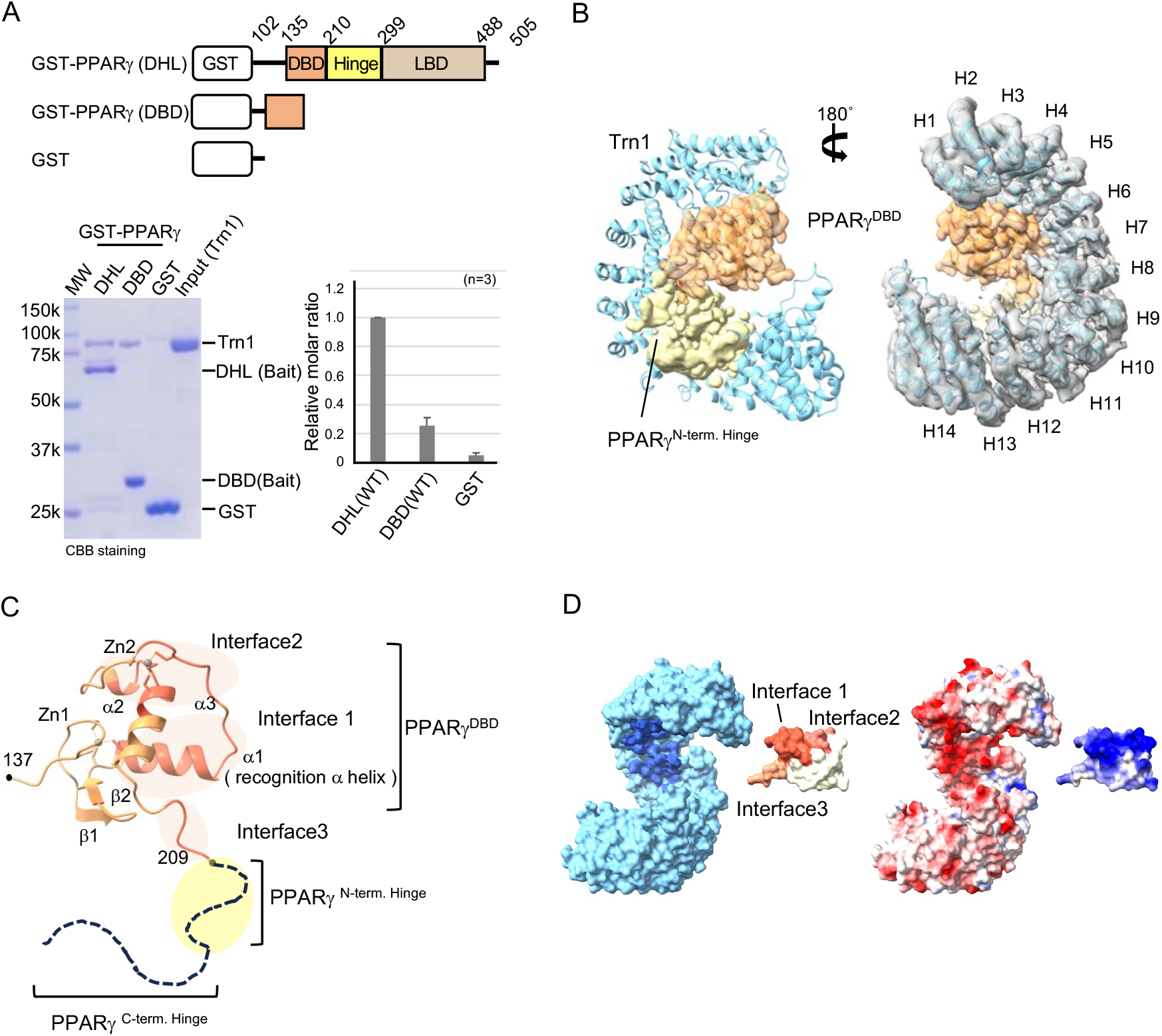
Cryo-EM structure of PPARg (DHL) in complex with Trn1 (A) GST-pulldown assay for GST-PPARg (DHL) and GST-PPARg (DBD) with Trn1(WT). The top diagrams show the domain architecture of PPARg used in this study. The SDS-PAGE gel image (lower left) and its quantitative analysis (lower right) are shown. The graph shows the relative binding affinity of each mutant normalized against that of DHL(WT). The three independent experiments were conducted. Data are mean ± s.d. (B) A ribbon diagram of the complex with PPARg(DHL) map. Trn1 is shown in ribbon representation (blue) and PPARg^DBD^ and PPARg^N-term, Hinge^, are shown with cryo-EM map (left) and the 180° rotation model with cryo-EM map of Trn1 (right). (C) A ribbon diagram of the model structure of PPARg(DBD). The regions consisting of interfaces 1, 2, 3, and Zn ions are colored deep pink, and gray, respectively. (D) Open-book view of the molecular surface (left) and electrostatic surfaces (right). The interaction residues of the Trn1 and three interfaces of PPARg (DBD) are colored deep blue, deep pink, light pink, and salmon pink, respectively. Red and blue colors on the electrostatic surfaces indicate the negatively charged and positively charged surfaces, respectively.

### Overall structure of Trn1-PPARg complex

We determined the single particle cryo-EM structure of the human Trn1Δloop - PPARγ(DHL) complex at 3.74 Å resolution (*SI Appendix*, Fig. S6). Trn1 is a superhelical protein with 20 HEAT repeat motifs. The last two HEAT repeat structures (H19-20) and the GGSGGSG linker were disordered in the complex structure (*SI Appendix*, Fig. S2). We observed the density of the DBD and about one-third of the N-terminal side of the hinge region in PPARγ (DHL) (Fig. 1), but could not observe the remaining densities corresponding to the C-terminal hinge region and LBD (Fig.1*B, C*). Of interest, the density which likely corresponds to the LBD domain was observed in the 2D structure classification in Cryo-EM analysis when we used the cross-linked samples, but LBD was not observed in the 3D structural model (*SI Appendix*, Fig. S6 and S7) probably due to the flexible orientation. We could only model the DBD region because the density map of the hinge region is poor. Trn1 grips DBD by HEAT repeats 2 to 8 that correspond to the RanGTP binding site in Trn1. The N-terminal hinge region extended from DBD interacts with HEAT repeats from 9 to 12 and 17 (*SI Appendix*, Fig. S1 and S2).

The interface between Trn1 and PPARγ(DBD) can be classified into three regions (interfaces 1 to 3). The total buried surface area is 1108.7 Å^2^ consisting of three interfaces in which each interface 1, 2, and 3 is 460.3 Å^2^, 408.3 Å^2^, and 240.1 Å^2^, respectively (Fig. 1*B* and 1*C*). Electrostatic potential surface map indicates that electrostatic interactions are dominant (Fig. 1*D*).

### Molecular recognition of PPARγ(DBD) by Trn1

The complex structure revealed that Trn1 grips the folded DBD. The overall structures of DBD in Trn1 complex and in RXR/ PPARγ/DNA(*19*) are essentially the same with the Ca r.m.s.d. deviation of 2.3 Å except for the C-terminal region. The C-terminal region of the DBD changes its conformation dynamically. R209, the C-terminal residue of the DBD, is 11.2Å apart compared with that in the RXR/ PPARγ/DNA complex (Fig. 2*A*), probably because Trn1 attracted the hinge region to its cargo binding sites.

**Figure 2.**
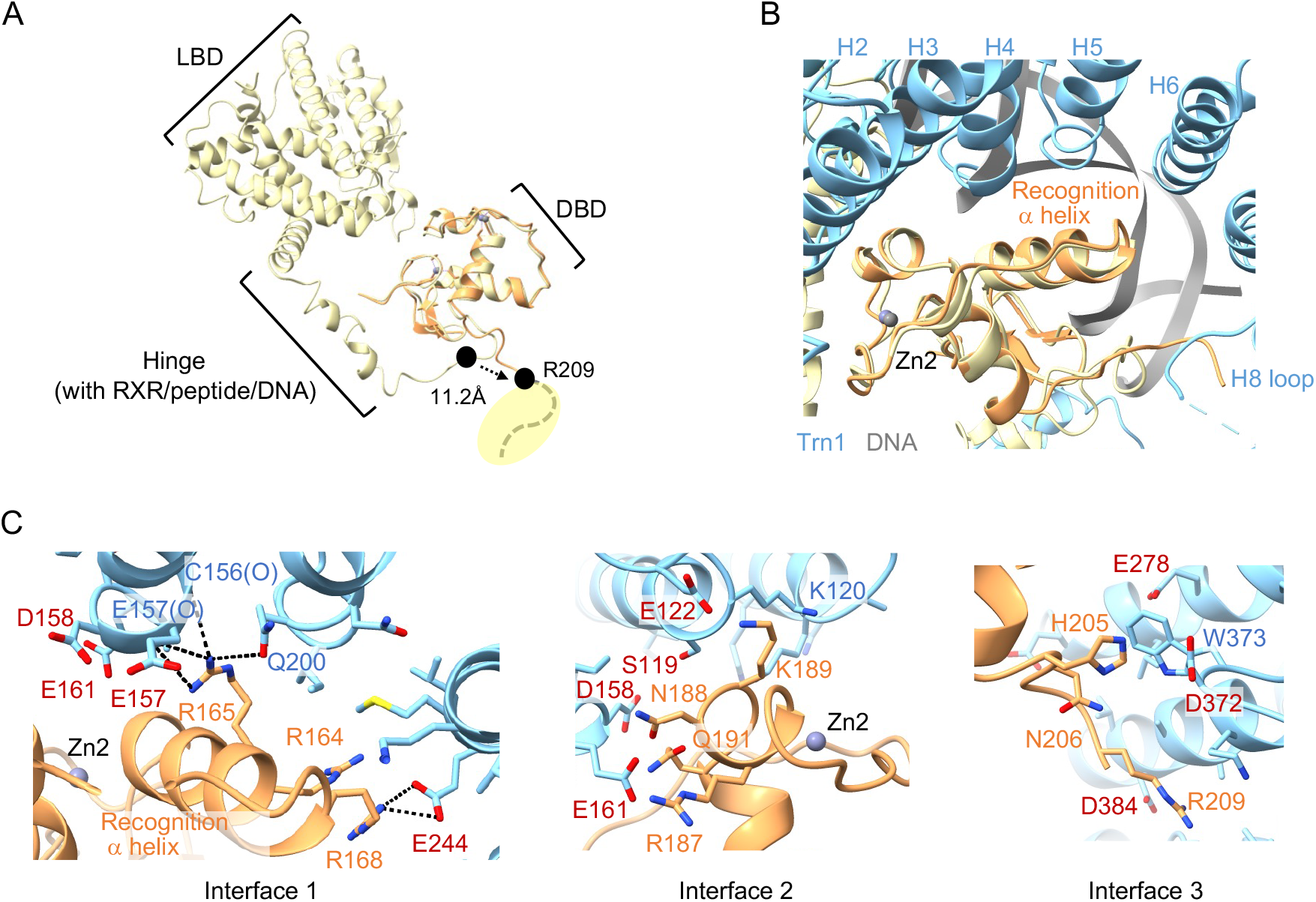
**I**nteraction details of PPARg in the complex (A) Superposition of Trn1/PPARg(DHL) complex structure colored in orange on the RXR/ PPARg/DNA complex structure colored in gold (PDB id: 3dzy). (B) Close-up view around recognition alpha helix. The gray model is the DNA in the RXR/ PPARg /DNA complex structure. (C) Close-up view of each interface. Stick side chain models indicate the amino acidic or basic acid residues of Trn1 and PPARg consisting of interfaces 1, 2, and 3.

The first a-helix, named as the recognition helix of DBD that inserts into the major groove of DNA in the RXR/PPARγ/DNA structure (*19*), is recognized by Trn1, making the dominant interaction in interface1 (Fig. 2*B*, *SI Appendix*, Fig. S8). There are basic residues on DBD and acidic residues on Trn1, which would contribute to the electrostatic interactions. In interface1, acidic residues of Trn1 (S159, S204, E157, D158, E161, E244, and E278) make contacts with the basic residues (R164, R165, and R168) of recognition a-helix. Among them, R165 makes multiple hydrogen bonds to the main chain oxygen atoms of C156 and E157 and oxygen of Q200. R168 makes a salt bridge to E244. In interface2, D158, S159, and E161 form an acidic cluster enabling to recognize R187, N188, K189A, and Q191. In interface 3, D372 exists near the H205, N206, and R209 (Fig. 2*C*, *SI Appendix*, Fig. S1).

### Amino acid residues essential for Trn1 binding and nuclear transportation of PPARγ

To investigate the essential residues to bind Trn1 and the transportation of PPARγ into the nucleus, we performed in vitro GST-pulldown and in-cell nuclear transportation assays. We mutated acidic residues of Trn1 and basic residues of PPARγ in interfaces 1, 2, and 3 to Ala to evaluate an influence on electrostatic interactions between them. Moreover, we made a mutant truncating 213-218 in the N-terminal hinge region to examine the effect of the binding although the density maps around the hinge region are poor. This mutant diminished the binding ability, suggesting that the region affects the binding to some degree. (Fig. 3*A*).

**Figure 3.**
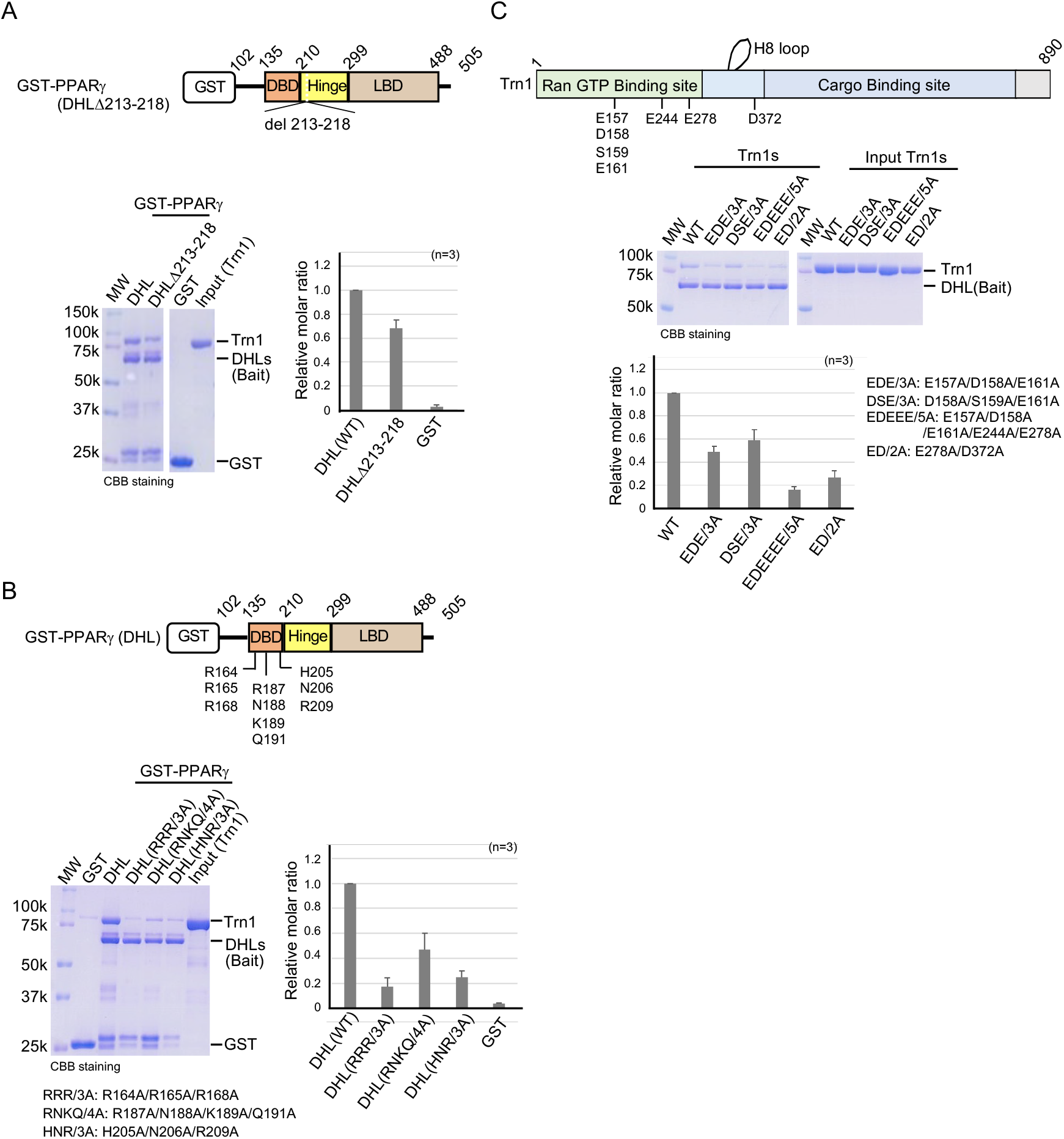
GST-pulldown assay of mutants (A) The result using GST-PPARg (DHLD213-218) mutant. (B) The result using GST-PPARg(DHL) Ala mutants. (C) The result using Trn1 mutants. The top diagrams show the domain architecture of PPARg or Tn1 used in the assay. The lower left photos are the result of SDS-PAGE analysis. The bar chart shows the relative average intensity normalized against the intensity of DHL(WT). n=3 independent experiment for all experiments, errors shown are s.d.

All triple/quadruple Ala mutants of PPARγ(DHL) significantly decreased the binding affinity to Trn1. R164A/R165A/R168A (RRR/3A) mutant on recognition a-helix on interface1 decreased the binding ability (Fig 3*B*). Consistently, all four mutants of Trn1 contributing to the interaction in interfaces 1, 2, and 3 also diminish the ability (Fig. 3*C*).

Next, to examine the effects of RRR/3A, RNKQ/4A, and HNR/3A mutations on the subcellular localization of PPARγ(DBD) and PPARγ (DHL), PPARγ fused with enhanced yellow fluorescent protein (EYFP) were expressed in HeLa cells (Fig. 4). EYFP, which has no localization signal, is diffused both in the cytoplasm and nucleus, while PPARγ (DBD WT)-EYFP and PPARγ (DHL WT)-EYFP are localized to the nucleus (Fig. 4*B, C*). The RRR/3A, RNKQ/4A, and HNR/3A mutations reduced the nuclear localization ratio of PPARγ (DBD)-EYFP, with RRR/3A exhibiting the most reduced effect. The mutations of PPARγ (DHL)-EYFP show a similar reduced localization ratio, but the effect is smaller than that for PPARγ (DBD)-EYFP, possibly due to the stronger affinity. These results suggest that the amino acid residues at the interface with Trn1 (R164, R165, R168, R187, N188, Q191, and R187, H205, N206, and R209) are essential for the nuclear localization of PPARγ.

**Figure 4.**
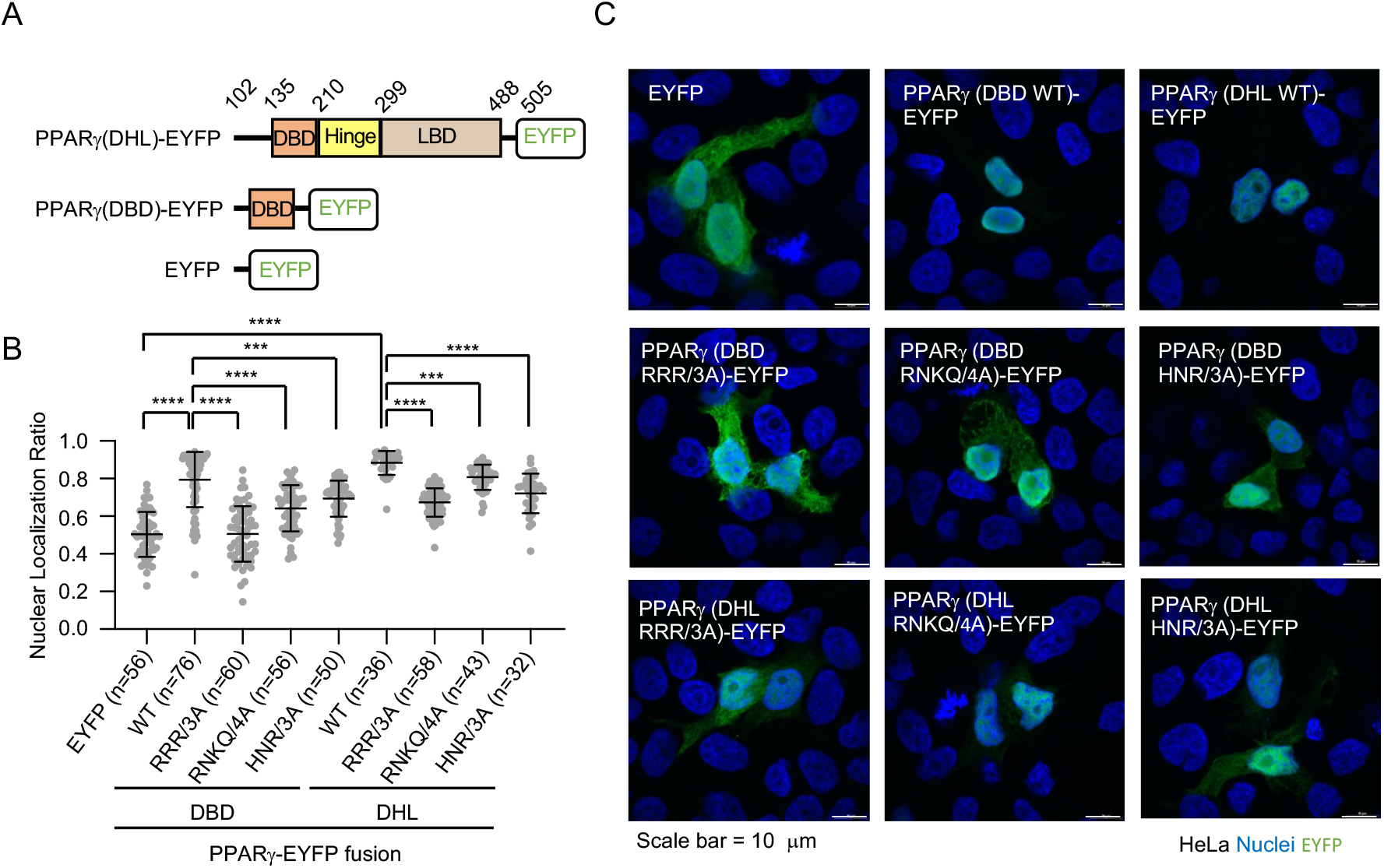
Nuclear localization assay (A) The domain architecture of the proteins used in the assay. (B) Subcellular localization of EYFP and PPARg-EYFP proteins in HeLa cells. Green: EYFP and PPARg-EYFP proteins. Blue: nuclei stained with DAPI. Bar = 10 μm. C, Plot of nuclear localization ratio of each protein. Numbers in parenthesess indicate the number of cells observed. Data shown are the mean ± s.d. Each point on the graph is the nuclear localization ratio of an individual cell. ***, p < 0.0005; ****, p < 0.0001.

### Structure comparison of Trn1s in complex with RanGTP or cargo peptides complexes

Next, we compare the recognition manner of RanGTP or cargo peptides by Trn1(*20, 21*). Trn1 enfolds all binding proteins/peptides using its concave surface. Trn1 uses a wide area for gripping the PPARγ covering the RanGTP binding site. The buried surface area of the RanGTP complex is 2056.8 Å^2^, which is almost twice the area of the DBD-binding surface of the PPARγ complex (1108.7Å^2^). Accordingly, RanGTP binds to Trn1 stronger than PPARγ. Actually, the *K*D of PPARγ (DHL) to Trn1 was 200 nM (*SI Appendix*, Fig. S3), and the reported binding affinity of RanGTP to Trn1 was hundreds of picomolar(*1, 22*), indicating that RanGTP and PPARg bindings to Trn1 are mutually exclusive, and RanGTP binding could release PPARγ efficiently.

Trn1 also grips the N-terminal hinge region using with HEAT repeats from 9 to 12 and 17, which is part of the classical cargo binding site of Trn1. Especially, site A is mainly occupied (*SI Appendix*, Fig. S9). The buried surface areas of the M9 peptide complex(*15*) and H3 peptide complex^13^ are 1760.2Å^2^ and 877.1Å^2^, respectively. The value of the DBD-binding surface (1023.1Å^2^) is slightly larger than that of the H3 peptide. (Fig. 5*A*, 5*B*, and *SI Appendix*, Fig. S9).

**Figure 5.**
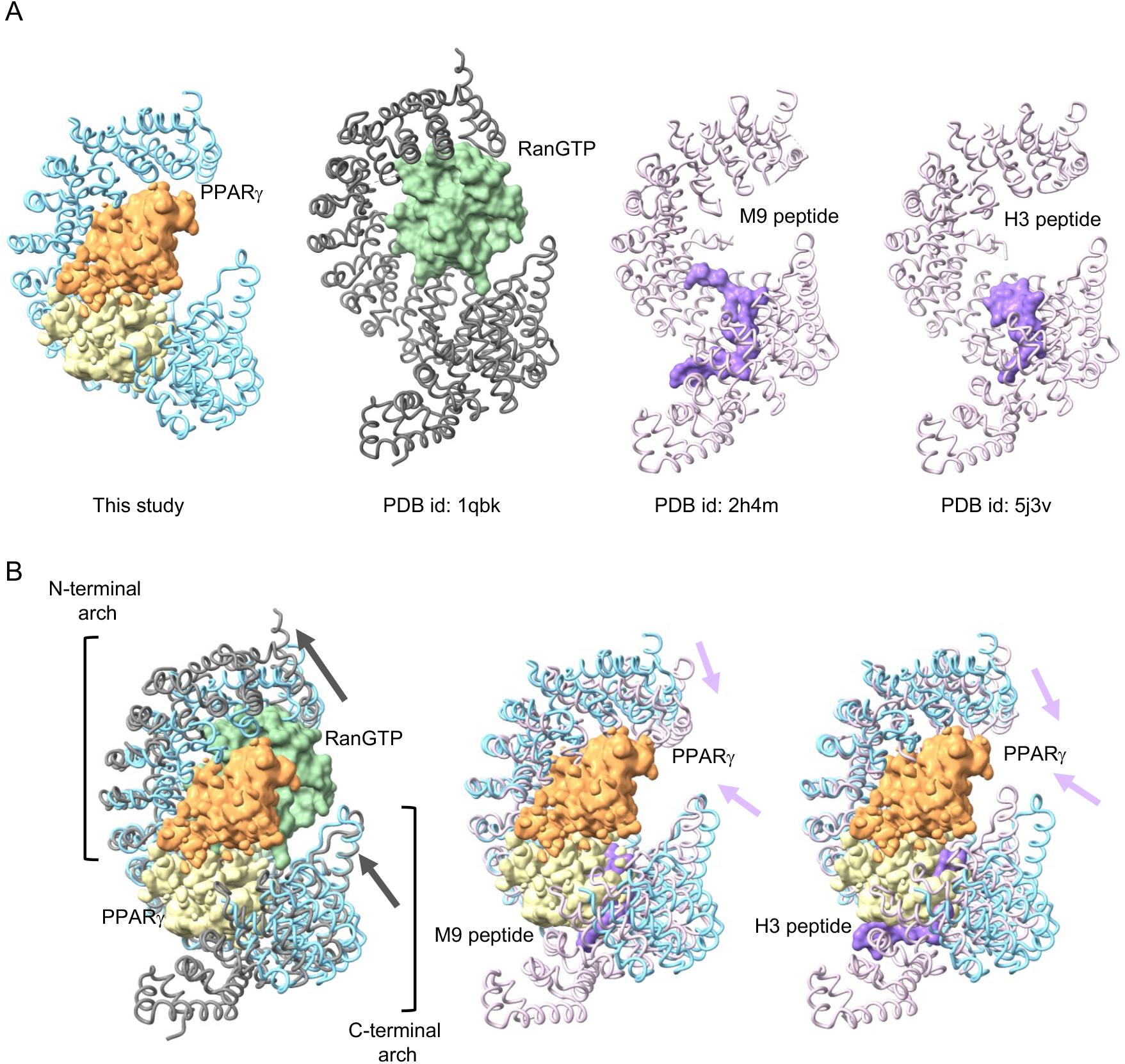
Structure comparison between PPARg/Trn1 and RanGTP/Trn1 or cargo peptide/Trn1 complexes (A) Trn1 complexes. The wire models showed Trn1 molecules in complex with PPARg (orange and yellow map), RanGTP (green surface), M9 peptide (purple surface), and H3 peptide (purple surface) colored in blue, dark gray, and purple, respectively. (B) The superposition of each Trn1 model.

Superposition of Trn1 in Trn1-PPARγ and Trn1-RanGTP complexes revealed that dynamical structural rearrangement occurs (Fig. 5). Only the central small region (265 pruned atom pairs mainly from HEAT repeats 8-14) of Trn1 is overlapped well with the RMSD 1.1 Å. However, both N- and C-terminal arches change their relative position to fit the binding protein, giving rise to the large RMSD of 6.5 Å for the whole protein (729 atom pairs). The RanGTP-binding pushes away the Heat repeat motifs in the N-terminal arch of Trn1 and attracts the Heat repeat motifs in the C-terminal one to the center of the complex structure. In contrast, the cargo peptide binding, M9 with PY-NLS or H3 without PY-NLS, attracts the Heat repeat motifs in both the N- and C-terminal arches. Trn1 in Trn1-PPARg is superimposed well around H5-H11 (282 pruned atom pairs with RMSD 1.0 Å) in the M9 complex and around H6-H13 (286 pruned atom pairs with RMSD 1.0 Å) in the H3 complex, but the RMSD values of the whole Trn1 are large with 6.0 Å (721 pairs) and 5.2 Å (732 pairs), respectively. Large structural rearrangement in Trn1 is induced for the various binding partners.

## Discussion

PPARs activate specific genes in a ligand-dependent manner, and nuclear transportation is one of the essential processes exerting their functions in the cell. High molecular weight proteins (more than ca. 40KDa) need escort by nuclear transportation receptors to enter the nucleus and harbor a nuclear localization signal to be caught by the receptors. PPARγ has no canonical NLS sequence and its structural-based nuclear translocation mechanism has remained elusive for a long time. Recently, our group reported that Trn1 transports PPARγ to the nucleus in a redox-dependent manner (*12*). We provided structural insight into the nuclear transportation mechanism of PPARγ for the first time.

Trn1 recognizes PY-NLS(*15, 16*), NLS peptide without PY(*13*), and RGG regions(*17, 18*) of cargo proteins, but PPARγ has no such classical NLSs. Notably, our structure shows that Trn1 grips the folded DBD with the hinge region using a broad range of interfaces of PPARγ. Generally, classical NLS has specific sequence motifs, not a domain structure, recognized by nuclear transportation receptors. Our structural study revealed that multivalent interaction consisting of several interfaces with the folded DBD domain and the disordered linear elements in the hinge region also work as a nuclear localization signal region. Several structural studies of importinβ; and IPO9 showed that these import receptors bind folded domains, SREBP-2, Snail, and H2A-H2B(*23-25*). Our study added new insights into cargo recognition. Previously two NLSs (NLS1, NLS2) of PPAR family were reported, and our determined NLS region includes NLS1. Of interest, the sequence of NLS1 is highly conserved among PPAR family(*26*). but that of NLS2 is not.

Trn1 has a long H8 loop that works as cargo dissociation or recognition(*18, 21*). The recently determined Cryo-EM structure of HNRNPH2 peptide in complex with full-length Trn1 with intact H8 loop showed that the H8 loop contains two short helices proximal to the helices of H8 and the middle region of the loop is disordered(*27*). The density map of PPARg does not clash the two helices (*SI Appendix*, Fig. S10), indicating that the H8 loop would have no inhibitory effect for complex formation with PPARγ.

The amino acid sequence of the DBD domain is highly conserved in the nuclear receptor family. The amino acids selected in the mutagenesis study on interface1 and interface2 are partially conserved and are variable on interface3 (*SI Appendix*, Fig. S11). The N-terminal hinge region is also a part of NLS. It is well known that the conserved DBD region has a function to recognize target DNA. Moreover, the short carboxy C-terminal extensions of DBD make an extensive DNA interaction with the upstream AAACT sequence of Peroxisome Proliferator Response Element (PPRE element) in the RXR/PPARγ/DNA complex structure(*19*). This extension corresponds to the N-terminal hinge region. Mutation analyses showed that both DBD and the hinge region cooperatively exert the nuclear localization ability of PPARγ, revealing that the hinge region has another function as a part of NLS in addition to the DNA binding. Conservation of amino acid sequence in DBD raises the possibility that Trn1 may be the predominant nuclear transporter for nuclear receptor family proteins. Among the amino acids selected for mutation analysis, missense mutations in the DBD have been reported to be associated with lipodystrophy subtype 3 (R164W, R165T) and cancer (R164W, R168K) (*28*). These missense mutations have been reported to reduce the expression of PPARγ -responsive genes(*29, 30*). These missense mutations in DBD may weaken not only the interaction between PPARγ and DNA but also that between Trn1 and PPARγ, thereby inhibiting PPARγ nuclear translocation. Contrary to DBD, the amino acid sequences of the hinge region are not conserved among the family (*SI Appendix*, Fig. S11). This region may function as a discriminator to Trn1.

Nuclear receptor family proteins harbor a sequence variable N-terminal domain and functional C-terminal domains/region (DBD, hinge region, and LBD). There are enormous structures of LBD (over 200) in the Protein Data bank. On the other hand, the reported atomic structure containing all functional regions (DBD-Hinge-LBD) is the only one in the RXR/PPARγ/DNA complex structure(*19*). This study is the second structural report of PPARγ containing the DBD-Hinge-LBD and is the first snapshot structure working in nuclear transportation, not in transcription. We found that the DBD-Hinge-LBD protein is unstable and degraded during purification, but the complex formation with Trn1 made the PPARγ more stable and reduced degradation. We confirmed that PPARγ also remained intact in the complex using Cryo-EM analysis but could not observe the density of the C-terminal region of the Hinge region and LBD domain, suggesting that LBD is not involved in the nuclear transportation process. LBD forms a dimer with other receptors. This result may give us an insight that Trn1 can transport heterodimerized receptors to the nucleus together. Recent studies reveal that Trn1 works as an inhibitor of liquid-liquid phase separation of FUS in addition to the nuclear transportation receptor(*31*). Our study demonstrated that Trn1 grips the folded DBD in the cell. This result raises the possibility that Trn1 may function in the transcription process of nuclear receptor family proteins, not only the function of nuclear transportation.

## Materials and Methods

### Expression of Trn1 and PPARγ(DBD) and PPARγ (DHL)

The full-length cDNA of human Trn1 subcloned into the plasmid vector pGEX6p-1 was given by Pro. Mamoru Sato (Yokokohama City University). The full-length cDNA of human PPARγ(NM_015869.5) was subcloned from the cDNA Library purchased from GenoStaff. Then, the regions of PPARγ (DHL:102-505) and PPARγ (DBD:135-209) were separately subcloned into the pGEX6p-1. The Δloop mutant of Trn1 was prepared by deleting amino acid 344-375 and then inserting the GGSGGSG sequence. Deletion- and point-mutant plasmids were generated using the QuikChange Lightning Site-Directed Mutagenesis Kit (STRATAGENE) or PrimeSTAR Mutagenesis Basal Kit (TaKaRa). We used E. coli BL21 (DE3)RIPL or Rosetta2(DE3) cells as host strains for protein expression. We followed the previously reported procedures of expression and purification for Trn1 and PPARg(*19, 21*). Finally, all proteins were purified by Superdex200pg (Cytiva) size exclusion chromatography with the buffer containing 110 mM CH3COOK, 10 mM DTT, 20 mM Hepes-KOH pH7.3.

### Preparation of Trn1/PPARγ complexes

For preparing Trn1/PPARγ (DHL), Trn1Δloop/PPARγ (DHL), and Trn1Δloop/PPARγ (DBD) complexes, both Trn1 or Trn1Δloop solution and PPARγ (DHL) or PPARγ (DBD) were mixed in molar ratio 1:1.5∼2.0 and incubated on ice. For preparing Trn1Δloop/PPARγ (DBD) complex, the 18 mg/mL PPARγ (DBD) solution was mixed with the Trn1Δloop solution in a molar ratio of 40:1. The white precipitations appeared and were then collected using a centrifugation (20000g, 10 min, 4°C). The pellet was dissolved with a buffer (110 mM CH3COOK, 20 mM Hepes-KOH pH7.3, 200 mM Arginine hydrochloride). Glutaraldehyde was used as a cross-linker to prepare the cross-linked Trn1Δloop/PPARγ (DHL) complex. The buffer was exchanged for a reaction solution (20 mM Hepes-KOH pH 7.3 or 8.2) before. Glutaraldehyde (Nacalai Tesque) was added to the complex solution (0.3 or 0.8 mg/mL) in the final concentration of 0.1%(v/v), and then the reaction solution was incubated for 30 min at 20°C. The reaction completion was confirmed by SDS-PAGE analysis. Each complex was purified by Superdex200 Increase 10/300 (Cytiva) size exclusion chromatography with the buffer containing 110 mM CH3COOK, 10 mM DTT, and 20 mM Hepes-KOH pH7.3. Complex formation was confirmed by SDS-PAGE analysis.

### ITC analysis

Isothermal titration calorimetry (ITC) was performed to measure the binding equilibrium between Trn1 and PPARγ (102-505) using MicroCal iTC200 (GE Healthcare) in a buffer (20 mM HEPES KOH pH 7.3, and 100 mM NaCl) at 298K. The data was analyzed by using OriginLab Software (GE Healthcare). The protein concentration of Trn1 (in Cell) and PPARγ (DHL) (in syringe) were 10 mM and 100 mM, respectively.

### Single particle cryoEM structure determination

Trn1Δloop/PPARγ (DHL) was used for structural determination. To avoid the destruction of the complex in the cryo-grid, the buffer dissolving the complex was exchanged for a buffer (10 mM DTT and 20 mM Hepes-KOH pH7.3.) using Superdex200 Increase 10/300 (Cytiva) size exclusion chromatography. Samples were frozen as soon as possible after elution and stored at -80°C until grid preparation. 2.6 µL of Trn1Δloop/PPARγ (DHL) solution was applied to a glow-discharged Quantifoil holey carbon grid (R1.2/1.3, Cu, 300 mech), blotted for 4.5 sec at 4ºC and plunge-frozen into liquid ethane using a Vitrobot Mark IV (Thermo Fishrer Scientific). The grid was inserted into a Titan Krios (Thermo Fishrer Scientific) operating at an acceleration voltage of 300 kV and equipped with a Cs corrector (CEOS, GmbH). Images were recorded with a K3 direct electron detector (Gatan) in Counting mode with an energy filter at a slit width of 20 eV. Data were automatically collected using SerialEM software (*32*) at a physical pixel size of 0.675 Å, with 64 frames at a dose of 0.94 e^-^/Å per frame, an exposure time of 3.196 sec per movie, and defocus ranging from -0.8 to -1.5 µm. A total of 7,778 movies were collected. The movie frames were subjected to beam-induced motion correction using MotionCorr2.1 (*33*), and the contrast transfer function (CTF) was evaluated using Gctf (*34*). Approximately 1,000 particles were manually selected from 10 micrographs to perform two-dimensional (2D) classification. Using a good 2D class average image as a template, a total of 6,256,331 particle images were automatically picked from all micrographs and were extracted with a box size of 72 pixels with 4x binning using RELION-3.1(*35*). After three rounds of 2D classification, 749,125 particles were selected and subjected to 2D classification and ab-initio using cryoSPARC2(*36*) for building the initial model of the Trn1Dloop/PPARg (DHL), as shown in *SI Appendix*, Fig. S6*D*. After ab-initio, 717,475 particles were re-extracted at a pixel size of 0.675 Å and subjected to two rounds of Heterogeneous Refinement. A total of 287,381 particles were subjected to Homogeneous Refinement and Non-uniform Refinement. The final 3D refinement and post-processing yielded a map with global resolution of 3.74 Å, according Fourier shell correlation (FSC) with the 0.143 criterion. Local resolution was estimated using cryoSPARC2. The processing workflow is outlined in *SI Appendix*, Fig. S6*A*. Structure modeling was performed using ChimeraX(*37*) and Coot(*38*). The models were refined in real space using Phenix(*39*) and validated for stereochemical and geometrical suitability with MolProbity(*40*). The details are summarized in *SI Appendix*, Table S1.

### Size Exclusion Chromatography/Multiple Angle Light Scattering analysis (SEC-MALS)

SEC-MALS analysis was performed to estimate the molecular weight of the complex. 20 uL of 6∼7 mg/mL of each sample was injected to superdex200 Increase 10/300 (Cytiva) column. The solution (110 mM CH3COOK, 20 mM Hepes-KOH pH7.3, 10 mM DTT) was used as the mobile phase, and the flow rate was 0.5 ml/min. Light scattering and differential refractive index were measured using a multi-angle light scattering DAWN HELEOS II detector (Wyatt Technology), and a 2414 Refractive Index Detector (Waters) respectively.

### SEC-Small angle X-ray Scattering analysis (SEC-SAXS)

SEC-SAXS experiment was performed on beamline BL-10C at PF (Tsukuba, Japan). 140∼150 uL of 6∼7 mg/mL of each complex solution was injected into the superdex200 increase 10/300 (GE Healthcare) column. SEC was performed with a buffer (110 mM CH3COOK, 20 mM Hepes-KOH pH7.3, 10 mM DTT) as the mobile phase. The flow rate was changed from 0.5 ml/min to 0.06 ml/min during protein elution. The A280 values monitored at the irradiation position were used for analysis. The program package SAngler(*41*) processed the scattering curves.

Solution structure was determined using the scattering data from the elution top peak. The radius of gyration (Rg) and the pair distance distribution functions (P(r) function were calculated using the program PRIMUS(*42*) and GNOM(*43*), respectively. The program DAMMIN(*44*) generated 15 dummy atom models independently. All models were averaged and selected using the program DAMAVER(*45*). Then the second DAMMIN calculation was performed using the damstart model derived by DAMAVER. The final model fitted the crystal structure. Superposition was performed with the program SUPCOMB(*46*). The details are summarized in *SI Appendix*, Table S2.

### Pulldown assay

GST-fused PPARγ(DHL) and PPARγ(DBD) proteins used in Fig. 3*A, B*, and *C* were expressed in 100 mL and 30 mL cultures, respectively. Harvested cells were lysed with Lysozyme and Triton-X. The cell lysate solutions were bound to 40 μL of GST-Accept resin (COSMOGEL). The resin was washed three times with 1 mL of wash buffer [50 mM Tris-Cl pH7.5, 300 mM NaCl, 10% glycerol, 2mM DTT, 100 uM Zinc Acetate, and 1x protease inhibitor cocktail (Nacalai Tesque)], and then washed with 1 mL of pulldown buffer (110mM CH3COOK, 20mM Hepes-KOH pH7.3, 10mM DTT). The protein-binding resins were incubated with 100 μg of purified full-length WT/mutants Trn1 for 0.5 h at 4°C, and then washed three times with 1 mL of the pulldown buffer, and eluted with 40 μL of 2x Laemmli’s sample buffer (Nacalai Tesque). The eluates were subjected to 12.5% SDS-PAGE (Nacalai Tesque) and stained with Coomassie Brilliant Blue (Fujifilm wako pure chemical). We used ChemiDoc XRS (Bio-rad) and ImageJ software (NIH) to take photos of SDS-PAGE gels and detect signals for quantification, respectively.

### Immunocytochemistry and image analysis

HeLa cells were cultured in Dulbecco’s Modified Eagle Medium (Nacalai Tesque) supplemented with 10% fetal bovine serum (Biowest) and an antibiotic–antimycotic solution (Nacalai Tesque) at 37°C in a 5% CO_2_ condition overnight. Transfection was performed using Fugene 6 (Promega). Transfected HeLa cells were washed with Dulbecco’s phosphate-buffered saline (PBS), fixed with 4% paraformaldehyde (PFA) dissolved in 100 mM phosphate buffer (PB, pH7.4) at room temperature (RT) for 10 min, washed with PBS, incubated in 10% normal donkey serum (EMD Millipore) and 0.2% Triton X-100 (Sigma-Aldrich) in PBS (hereafter referred to as the blocking buffer), incubated with rat anti-EGFP antibody(*47*) diluted in the blocking buffer at RT for 1 h, washed with PBS three times, incubated with donkey anti-rat IgG conjugated with Alexa Fluor 488 (Abcam) and DAPI (Dojindo) in the blocking buffer at RT for 1 h, washed with PBS three times, and mounted with Aqua-Poly/Mount (Polysciences). Confocal microscopy images were acquired using an AX (Nikon). Nuclear localization of the immunosignal was quantified using ImageJ software (NIH). The means of three or more groups were compared with one-way analysis of variance (ANOVA) with Dunnett’s multiple comparisons test using Prism 10 (GraphPad).

## Acknowledgments

We thank all beamline staff at BL44XU and BL32XU of SPring-8, BL-1A, BL-17A, and BL-10C of PF/KEK for their technical help and kind suggestions. We thank M. Sato for providing the plasmids and Kenji Iwasaki for supporting a Cyo-EM experiment. We also thank Miwa Kuzuhara and Akiko Ikegami for sample preparation.

This work was supported by a Grant-in-Aid from the Japanese Ministry of Education, Culture, Sports, Science, and Technology [S.T.F.(21K06031)and T.S.], AMED-CREST (S.T.F.) under Grant Number 22gm1410010s0202, CREST JST (T.S), and the Research Foundation for Pharmaceutical Sciences(S.T.F.), Platform Project for Supporting Drug Discovery and Life Science Research [Basis for Supporting Innovative Drug Discovery and Life Science Research (BINDS)] from AMED under Grant Numbers JP21am0101070, JP21am0101071, and JP21am0101072 (support number 0153 and 1587) and Takeda Science Foundation (S.O.).

