## Supplemental files for "Structural insight into the nuclear transportation mechanism of PPARγ by Transportin-1"

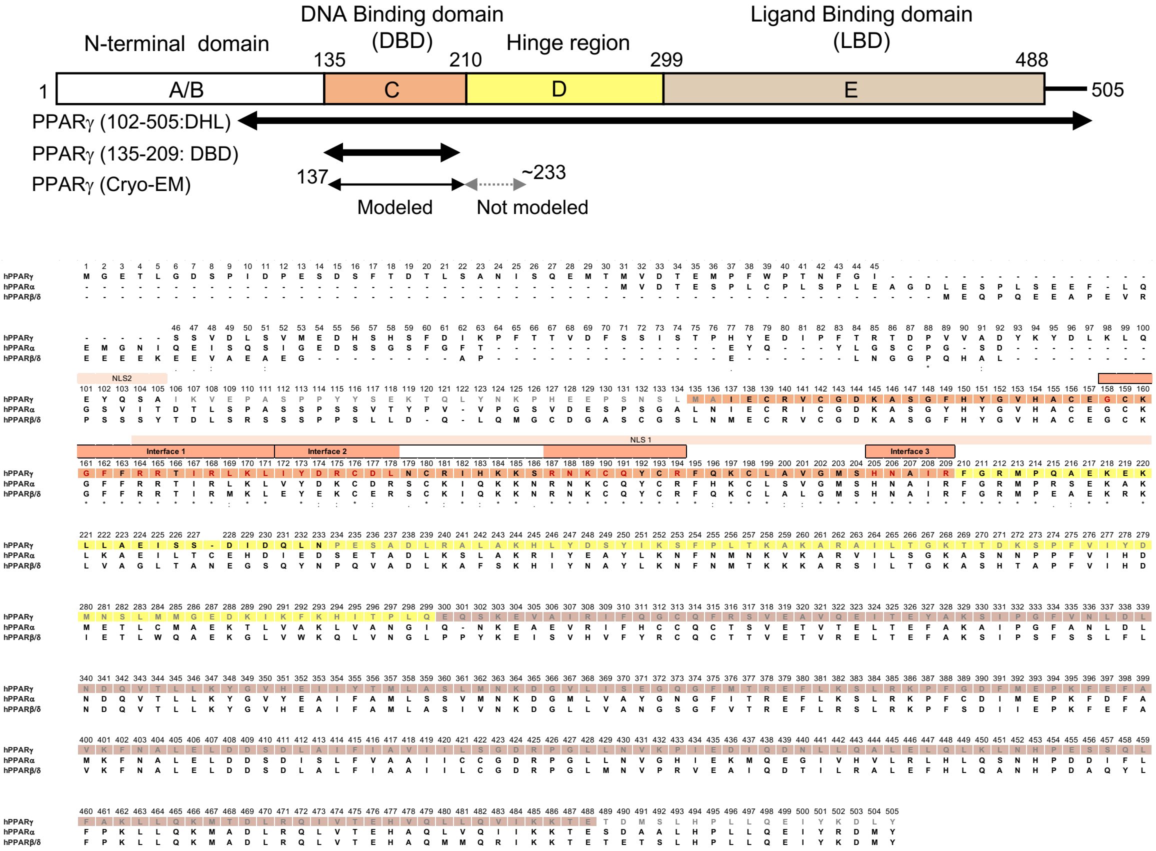


**Fig. S1.** Domain architecture and amino acid sequence of PPARγ. The upper diagram indicates the domain definition and the structure-determined region in this study.The amino acid sequences of PPAR family proteins are shown. NLS1 and NLS2 highlighted with gray color indicate the reported NLS sequence of PPARg[26]. The Upper boxes on the sequence indicate each interface. The highlight colors on the amino acid sequence of PPARγ are the same in the domain definition. Red characters indicate the interaction residues to Trn1.

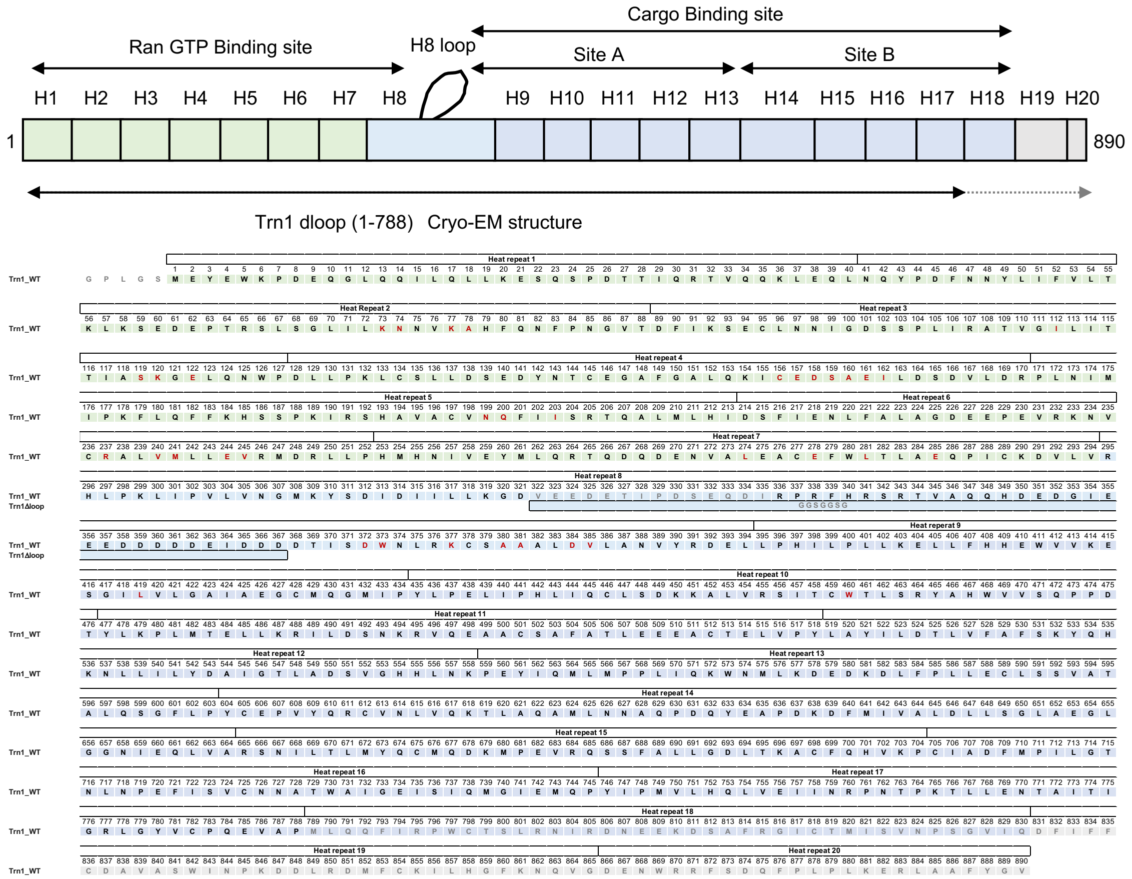


**Fig. S2.** Domain architecture and amino acid sequence of Trn1. The upper diagram indicates the definition of the binding sites and the structure-determined region in this study. The amino acid sequences of Trn1 and Trn1Dloop are shown. The Upper boxes on the sequence indicate each heat repeat. The highlight colors on the amino acid sequence of PPARγ are the same colors in the binding site definition. Red characters indicate the interaction residues to PPARγ.

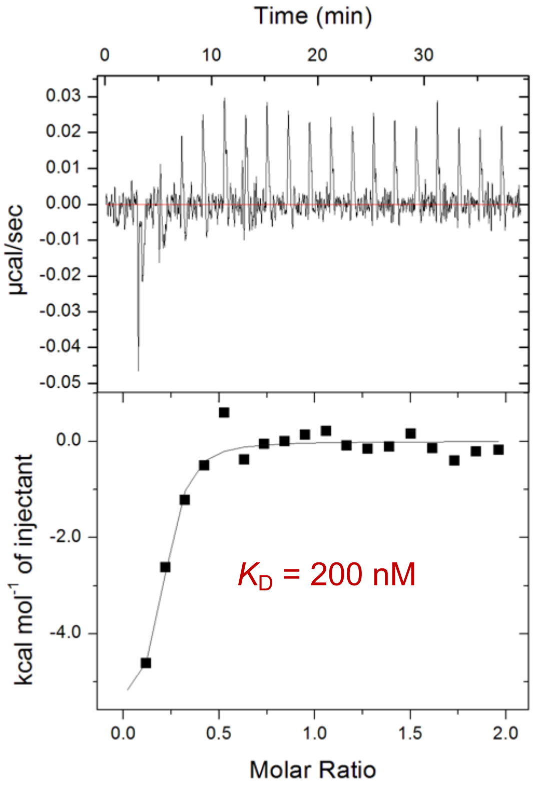


10 μM Trn1 titrated by 100 μM PPARγ (DHL)

**Fig. S3.** The results of ITC analysis**.** The titration plot(upper) and fitting curve (lower) are described. The derived dissociation constant ($K_D$) was indicated in red character.

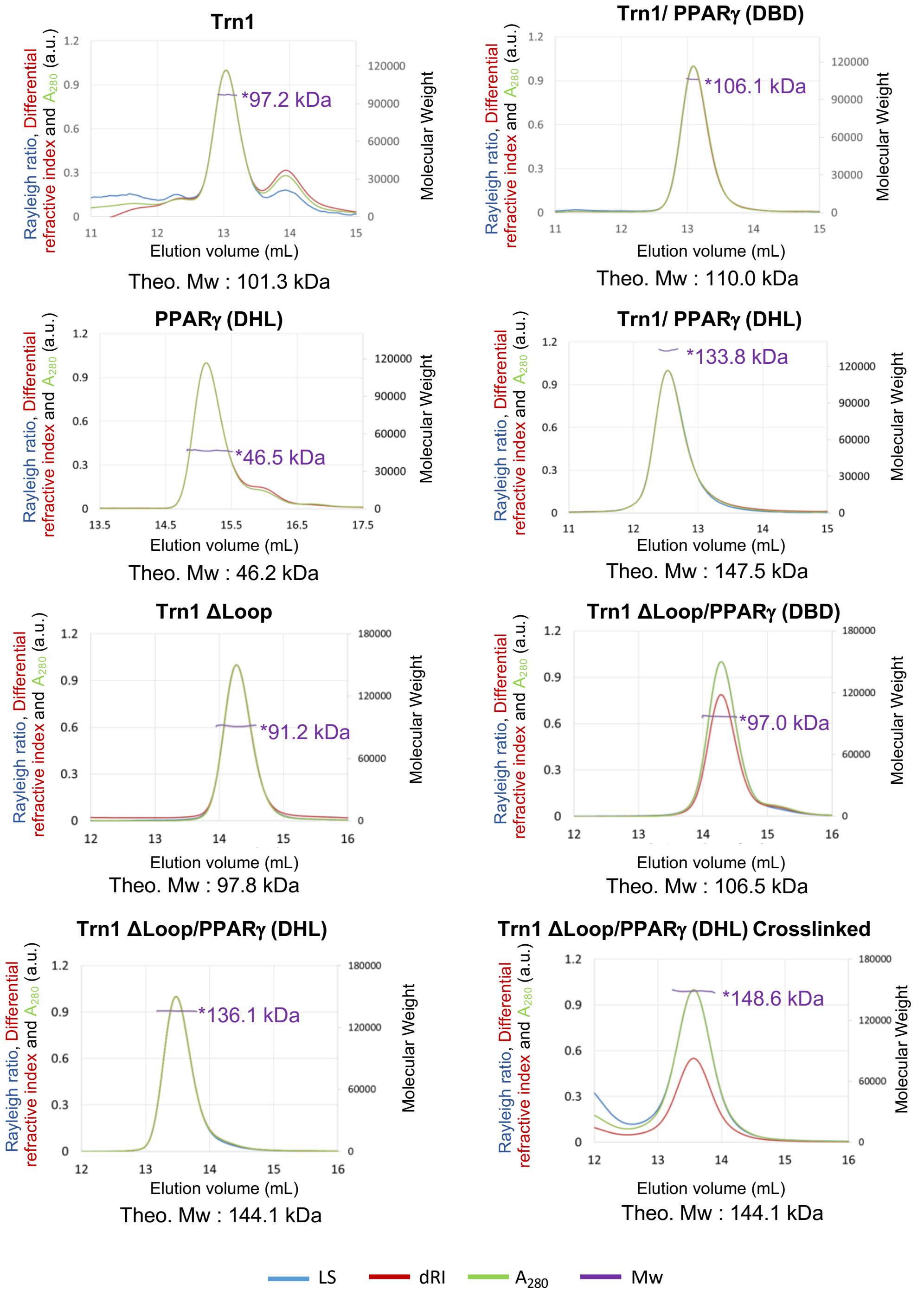


**Fig. S4.** The results of SEC-MALS analysis. Rayleigh ratio (LS: blue line), Differential refractive index (dRI: red line), $A_{280}$ (green line), and Molecular weight (Mw: purple line) are shown.

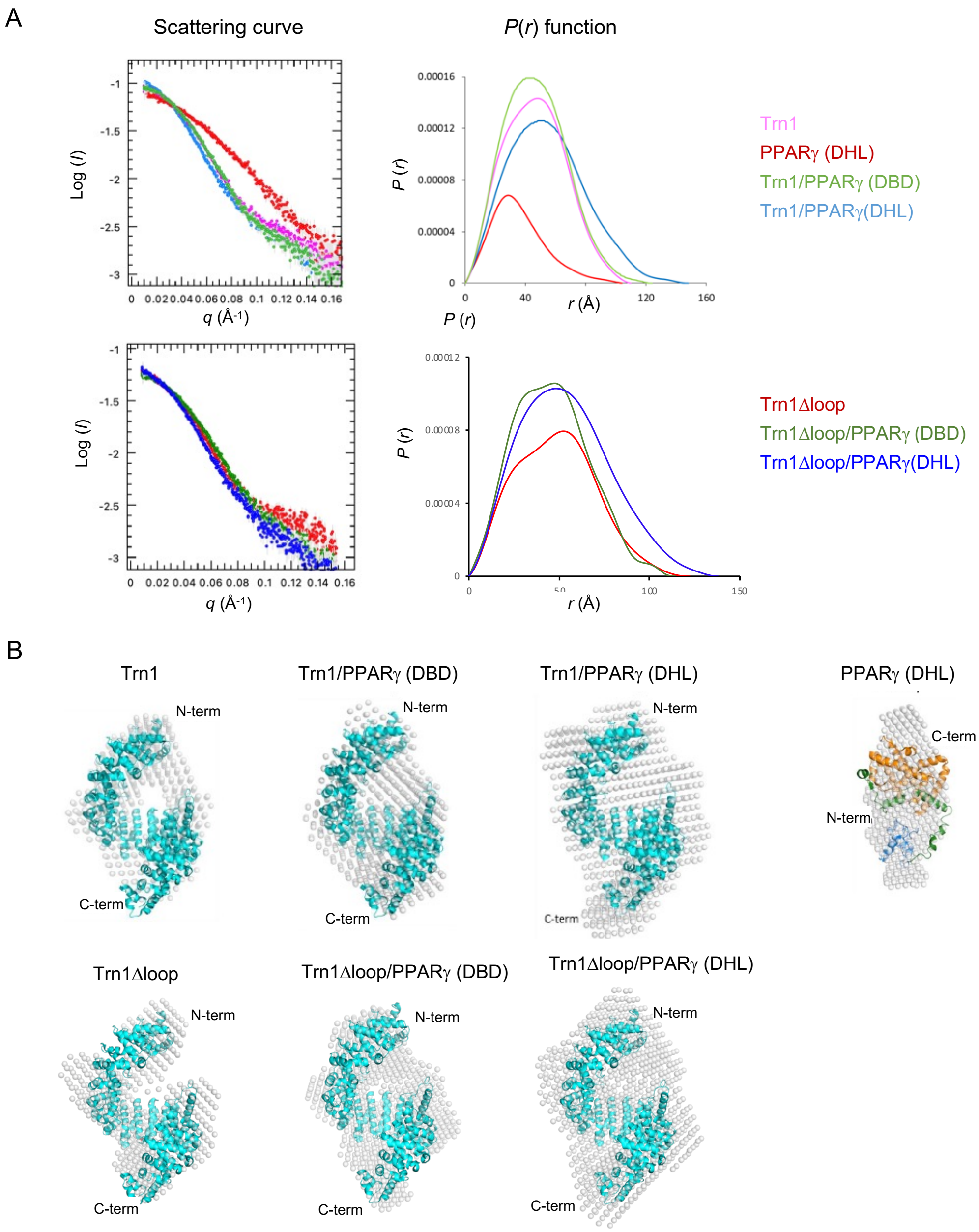


**Fig. S5.** Results of SEC-SAXS analysis. (A) Scattering curves (left) and P(r) function (right). (Upper) Pink, red, green, and sky blue lines indicate the plots of Trn1, PPARγ (DHL), Trn1/PPARγ (DBD), and Trn1/PPARγ (DHL), respectively. (Lower) red, green, and blue lines indicate the plots of Trn1Dloop, Trn1Dloop/PPARγ (DBD), and Trn1Dloop/PPARγ(DHL), respectively. (B) SAXS beads models. Blue Ribbon models are Trn1 structure (PDBid: 8Y70) superposed onto the beads models. The Ribbon model of PPARγ(DHL) extracted from PPARγ(DHL)/RXR/peptide/DNA complex (PDB id: 3dzy) fit into the beads model. Blue, green, and orange regions indicate DBD, Hinge, and LBD, respectively.

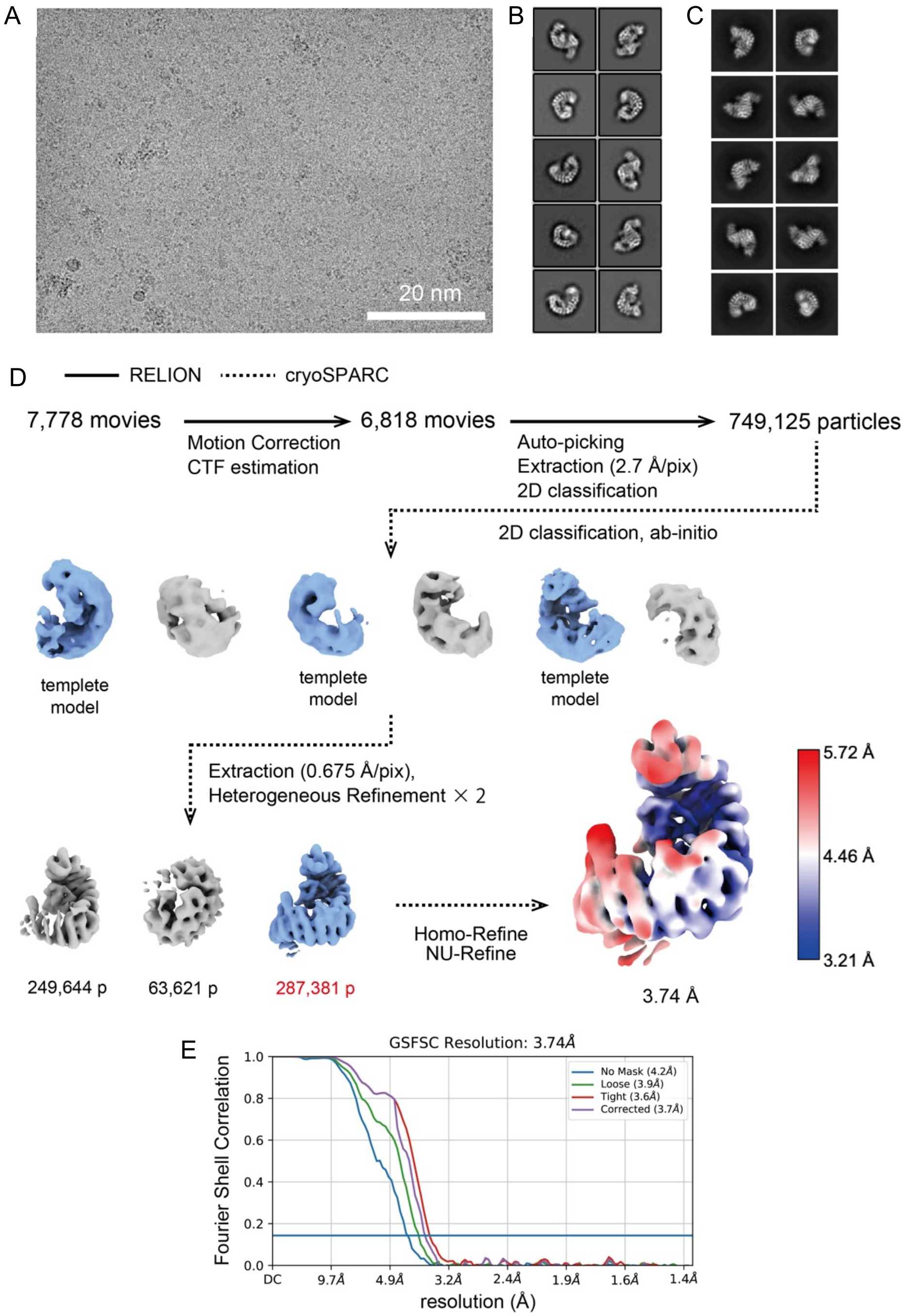


**Fig. S6.** Cryo-EM data processing. (A) A micrograph image of Trn1Dloop/PPARg (DHL). (B) 2D classification images of Trn1Dloop/PPARg (DHL) using RELION. (C) 2D classification images of Trn1Dloop/PPARg (DHL) using cryoSPARC2. (D) Cryo-EM processing workflow of Trn1Dloop/PPARg (DHL), displaying with the local resolution of EM density map. (E) Gold-standard unmasked and masked FSC curves using the 0.143 cutoff.

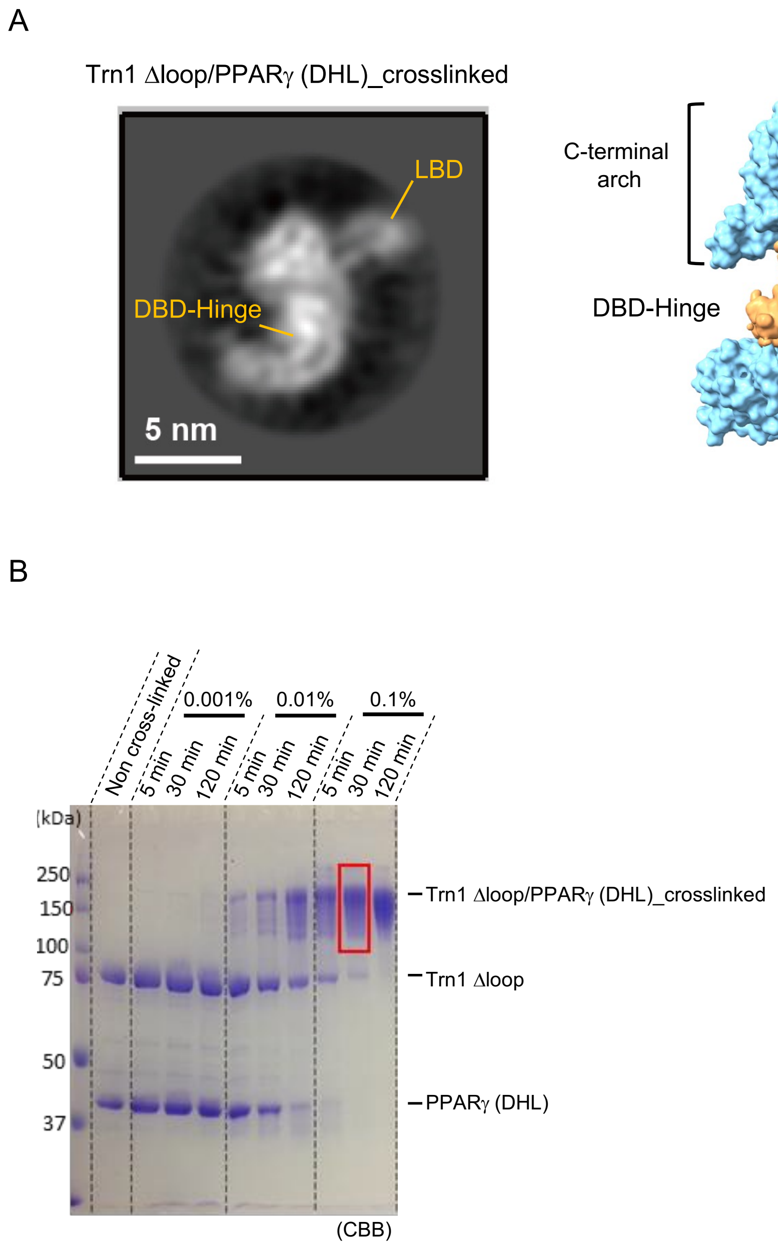


**Fig. S7.** 2D classification image of a crosslinked sample. (A) A 2D classification image of Trn1Dloop/PPARg (DHL)_crosslinked sample (left). A model structure corresponding to PPARg(DHL) (right). (B) SDS-PAGE analysis of cross-link samples. The condition highlighted by Red Box was used for sample preparation of Cryo-EM analysis.

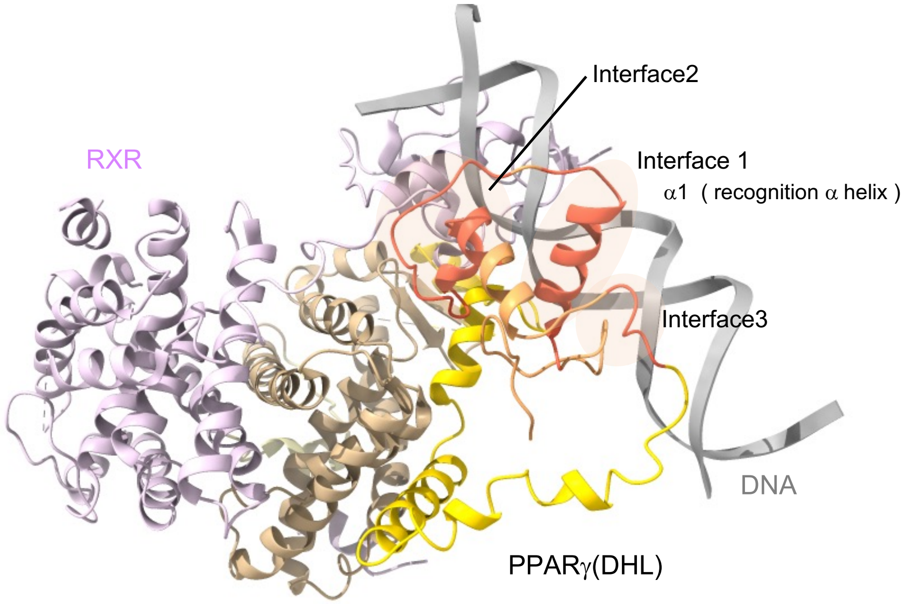


PDB id: 3dzy

**Fig. S8.** The mapping of the interfaces on RXR/PPARγ/DNA structure. The mapping of interfaces identified in the Trn1 complex on the /RXR/PPARγ/DNA complex (PDB id: 3dzy). All three interfaces were drawn in the same color drawn in Fig.1C

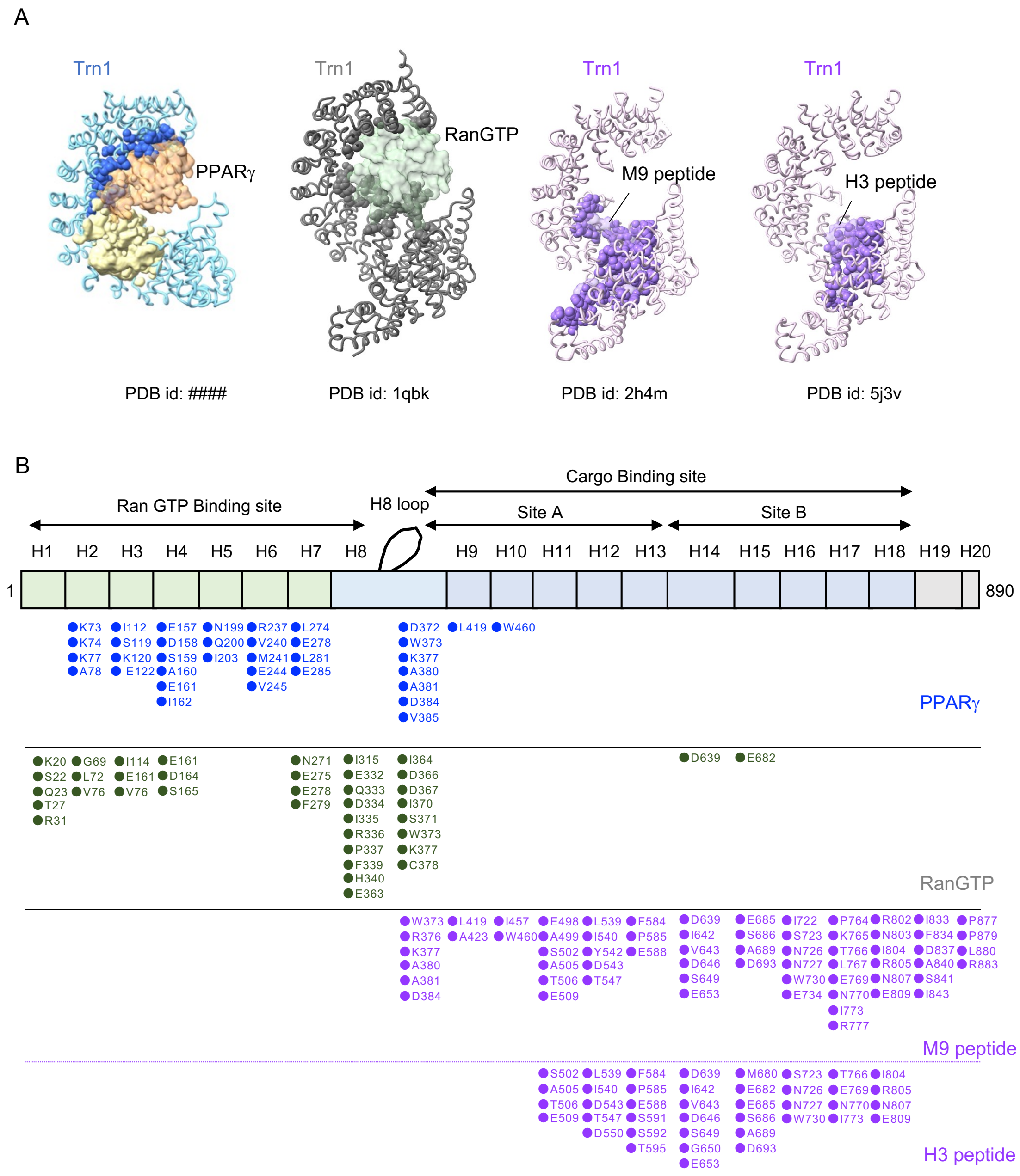


**Fig. S9.** Mapping of the interaction residues. (A) Diagrams show the Trn1 structures in complex with binding proteins [ PPARγ (PDB id: 8Y70) and RanGTP (PDB id: 1qbk)] and peptides [ M9 (PDB id: 2h4m) and H3 (PDB id: 5j3v). Tube models show the Trn1 on which the Sphere models indicate the residues involving the interactions. The transparent density (PPARγ)/surface models indicate the binding proteins or peptides. (B) The Interaction residues of Trn1 corresponding to sphere models in panel

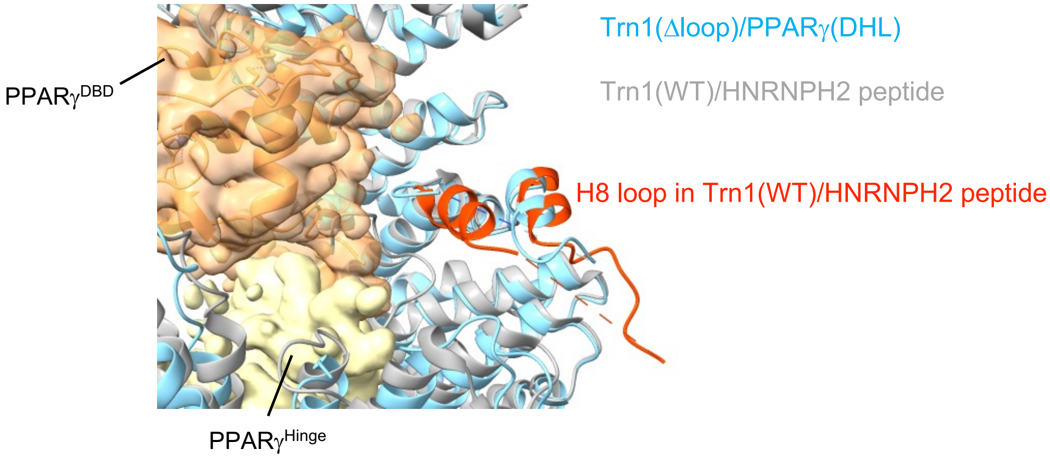


**Fig. S10.** Comparison of cryo-EM structure of Trn1(Δloop)/PPARγ(DHL) with Trn1(WT)/HNRNPH2 peptide. Cryo-EM structure of Trn1(Δloop)/PPARγ (DHL) with PPARγ (DHL) map was drawn in the same color in Fig.1B. Gray and orange ribbon diagram indicates Trn1 in Trn1(WT)/HNRNPH2 and H8 loop of the complex.

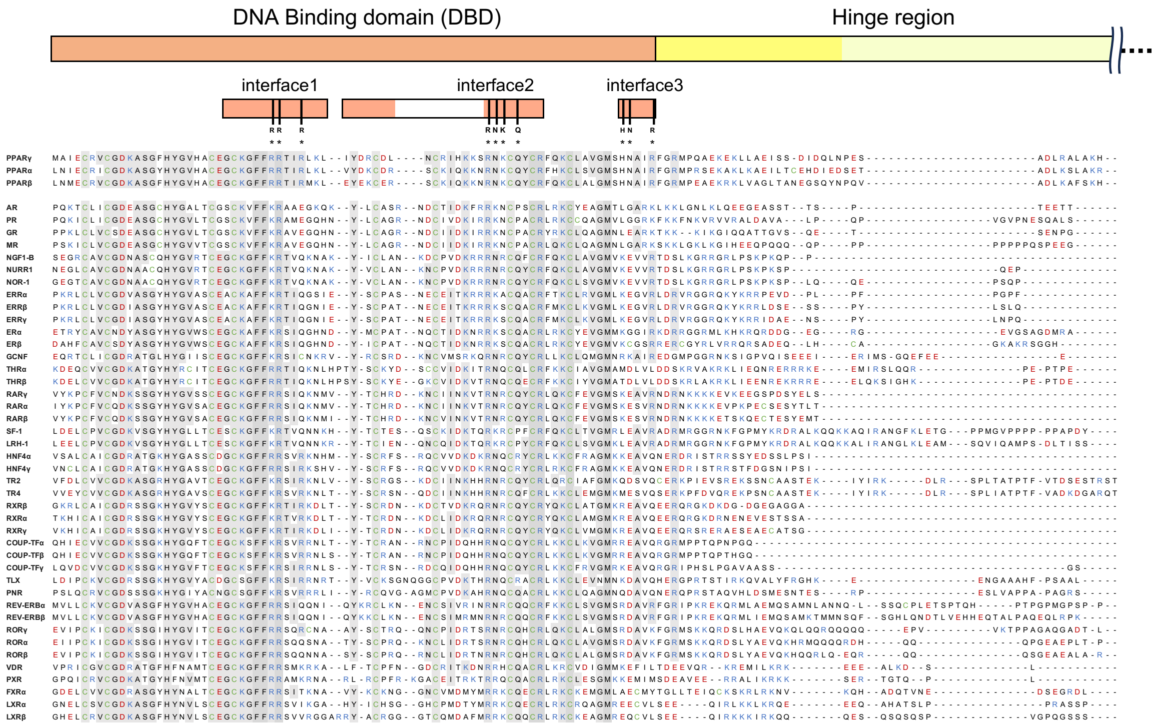


**Fig. S11.** Amino acid sequence alignment of nuclear receptor superfamily proteins. Sequence alignment of the DBD domain and N-terminal Hinge region of nuclear receptors. Red and Blue characters indicate acidic and basic residues, respectively. Green characters are Cys residues. Fully conserved residues are highlighted with gray boxes. Amino acid residues replaced by Ala in mutation analysis, which are RRR, RNKQ, and HNR on interface1, 2, and 3 are indicated.

**Table S1.** Cryo-EM data collection, refinement, and validation

| | Trn1Dloop/PPARg (DHL)<br>(EMD-39008)<br>(PDB 8Y70) |
|---|---|
| Data collection and processing | |
| Magnification | 105,000 |
| Voltage (kV) | 300 |
| Electron exposure (e–/Å$^2$) | 60 |
| Defocus range (μm) | -0.8 to -1.5 |
| Pixel size (Å) | 0.675 |
| Symmetry imposed | C1 |
| Initial particle images (no.) | 6,256,331 |
| Final particle images (no.) | 287,381 |
| Map resolution (Å) | 3.74 |
| FSC threshold | 0.143 |
| Map resolution range (Å) | 3.21-5.72 |
| Refinement | |
| Initial model used (PDB code) | 3dzy, 5j3v |
| Model composition | |
| Non-hydrogen atoms | 6389 |
| Protein residues | 806 |
| Ligands | 2 |
| B factors (Å$^2$)<br>(min/max/mean) | |
| Protein | 51.82/294.67/147.41 |
| Ligand | 159.67/214.17/186.92 |
| R.m.s. deviations | |
| Bond lengths (Å) | 0.003 |
| Bond angles (°) | 0.759 |
| Validation | |
| MolProbity score | 1.93 |
| Clashscore | 13.68 |
| Poor rotamers (%) | 0.42 |
| Ramachandran plot | |
| Favored (%) | 95.88 |
| Allowed (%) | 4.12 |
| Disallowed (%) | 0.00 |

**Table S2.** SEC-SAXS analyses

| Data-collection parameters | Trn1 | PPARg$^{(DHL)}$ | Trn1/PPARγ$^{DBD}$ | Trn1/PPARg $^{(DHL)}$ |
|---|---|---|---|---|
| Instrument | BL10C-SEC-SAXS (PF, Tsukuba) | BL10C-SEC-SAXS (PF, Tsukuba) | BL10C-SEC-SAXS (PF, Tsukuba) | BL10C-SEC-SAXS (PF, Tsukuba) |
| Wavelength (Å) | 1.00000 | 1.00000 | 1.00000 | 1.00000 |
| q range (Å$^{-1}$)[a] | 0.0106-0.2240 | 0.0134-0.2910 | 0.0128-0.2224 | 0.0107-0.1896 |
| Exposure time (sec.) | 20×1 frame | 20×1 frame | 20×1 frame | 20×1 frame |
| Temperature (K) | 293 | 293 | 293 | 293 |
| Structural parameters | | | | |
| I(0) (cm$^{-1}$) [from P(r)] | 0.10 ± 0.00 | 0.03 ± 0.00 | 0.11± 0.00 | 0.11 ± 0.00 |
| Rg (Å) [from P(r)] | 36.0 ± 0.15 | 28.5 ± 0.30 | 36.3 ± 0.26 | 42.6 ± 0.24 |
| I(0) (cm$^{-1}$) [from Guinier] | 0.10 ± 0.00 | 0.03 ± 0.00 | 0.11± 0.00 | 0.11 ± 0.00 |
| Rg (Å) [from Guinier] | 35.6 ± 2.41 | 27.5 ± 2.31 | 35.9 ± 3.36 | 42.1 ± 2.98 |
| Dmax (Å) | 111 | 104 | 124 | 148 |
| Porod volume estimate (Å$^3$) | 150062 | 77916 | 169684 | 244471 |
| M.W calculated with porod volume[b] | 93789 | 48698 | 106053 | 152794 |
| Dry volume calculated from sequence | 101300 | 46200 | 110000 | 147500 |
| Mean value of NSD (Dummy atom model) | 0.687 ± 0.019 | 0.539 ± 0.013 | 0.649 ± 0.013 | 0.675 ± 0.017 |
| Software employed | | | | |
| Primary data reduction | SAngler | SAngler | SAngler | SAngler |
| Data processing | PRIMUSQT | PRIMUSQT | PRIMUSQT | PRIMUSQT |
| Ab initio analysis | DAMMIN | DAMMIN | DAMMIN | DAMMIN |
| Validation and averaging | DAMAVER | DAMAVER | DAMAVER | DAMAVER |
| Computation of model intensities | CRYSOL | CRYSOL | CRYSOL | CRYSOL |
| Three-dimentional graphics representation | PyMOL | PyMOL | PyMOL | PyMOL |
